# PCP components control anterior and posterior regeneration, with a Prickle homolog impacting muscle organization, in the acoel *Hofstenia miamia*

**DOI:** 10.1101/2023.10.03.560791

**Authors:** D. Marcela Bolaños, Amber Rock, Núria Ros-Rocher, James Sikes, Mansi Srivastava

## Abstract

Whole-body regeneration requires wound response signals to control patterning programs to enable replacement of structures in their correct locations. While a number of molecular mechanisms underlying anterior-posterior regeneration have been identified, how small fragments of animals first re-establish polarity is less well understood, with non-canonical Wnt signaling recently emerging as a potential regulator. Here, we used the acoel worm *Hofstenia miamia*, a new research organism capable of robust whole-body regeneration, to assess functions of the components of the Planar Cell Polarity (PCP) pathway in establishing regeneration polarity. We identified homologs of Prickle (*pk-1*) and Diego (*dgo-1*) to be required for head and tail regeneration, respectively. RNA-sequencing analysis and experimental corroboration revealed that *pk-1* RNAi resulted in diminished expression of early wound response genes as well as of wound-induced expression of the anterior-specific marker *fz-7*, specifically in tail fragments. In contrast, *dgo-1* RNAi impacted wound-induced expression of the posterior-specific marker *tf7l2*, specifically in head fragments. Furthermore, *pk-1* and *dgo-1* are enriched in longitudinal muscle, with muscle fibers showing disorganized morphology at anterior-facing wound sites of tail fragments under *pk-1* RNAi. These findings suggest that *pk-1* and *dgo-1* are needed for wound-induced expression of anterior- and posterior-specific genes, and raise the possibility that this action is mediated via the control of muscle fiber orientation. Our work expands the study of PCP genes by revealing their functions in the process of whole-body regeneration in acoels, the sister-group to all other animals with bilateral symmetry, and will enable future studies of PCP components in controlling cellular and tissue-wide regeneration polarity.

## INTRODUCTION

Whole-body regeneration (WBR), the ability to fully restore any missing tissues and organs after injury, is widely observed across diverse animal phyla. This regenerative capacity requires precise control of the shape, size, and identity of newly formed tissue, which ultimately results in restoration of fully-functional organs (Blanchoud and Galliot, 2022; Srivastava, 2021). This precision is particularly evident in the reestablishment and patterning of the primary body axis upon transverse amputation, where small fragments of an animal regenerate heads and tails in the correct axial location. The question of how an animal knows to regenerate heads and tails in the right place has been heavily investigated, but the full decision-making that governs regeneration polarity is not known in any species.

Studies in three major model systems for WBR, cnidarians, planarians, and acoels, have implicated Wnt signaling as a key player in head and tail regeneration. Canonical Wnt signaling is central to the identity of new tissue along the primary body axis: inhibition of this pathway results in regeneration of heads instead of tails at posterior-facing wound sites in acoels and planarians, whereas excess of Wnt ligands promotes the regeneration of ectopic heads along the oral-aboral axis in *Hydra* (Chera et al., 2009; Gurley et al., 2010; Petersen and Reddien, 2009; Ramirez et al., 2020; Srivastava et al., 2014; Vogg et al., 2019). However, the control of polarity, i.e., the earliest decision upstream of Wnt activation in wounds that determines whether head or tail tissue should be made, is less well-understood. Recent data from planarians suggest that the earliest indicator of polarity, the asymmetrically expressed Wnt inhibitor *notum*, may be under the control of non-canonical Wnt signaling acting within muscle cells prior to injury (Gittin and Petersen, 2022). However, the specifics of this non-canonical Wnt signaling are unknown; for example, it is not clear whether this function is mediated with Wnt/PCP or Wnt/Ca+ pathways. Pre-injury inputs into early indicators of polarity are not well-studied in cnidarians and acoels.

The acoel worm *Hofstenia miamia,* is an emerging model organism that enables mechanistic studies of WBR (Gehrke et al., 2019; Hulett et al., 2020, 2023a; Ramirez et al., 2020; Srivastava, 2021). Given its capacity to fully regenerate heads and tails and its evolutionary divergence from other regenerative model invertebrates such as planarians (550 mya) and *Hydra* (650 mya), *Hofstenia* is a powerful system for investigating the mechanisms and evolution underlying regeneration polarity (Fig.1A). In *Hofstenia*, the earliest known activated genes that are asymmetrically detected in response to wounding are *wnt-3* and *sp5*, both upregulated at posterior-facing wounds (Ramirez et al., 2020). The pre-injury expression of a Wnt ligand, *wnt-1*, is needed for *wnt-3* and *sp5* expression, however it is uncertain whether this Wnt ligand is required for polarity of *wnt-3* and *sp5* expression and whether it acts through a canonical or non-canonical pathway. We therefore sought to study the roles of non-canonical Wnt signaling during regeneration in *Hofstenia*. Specifically, we assessed whether Wnt/PCP-mediated cellular polarity could contribute to tissue-wide polarity during regeneration.

**Figure 1.**
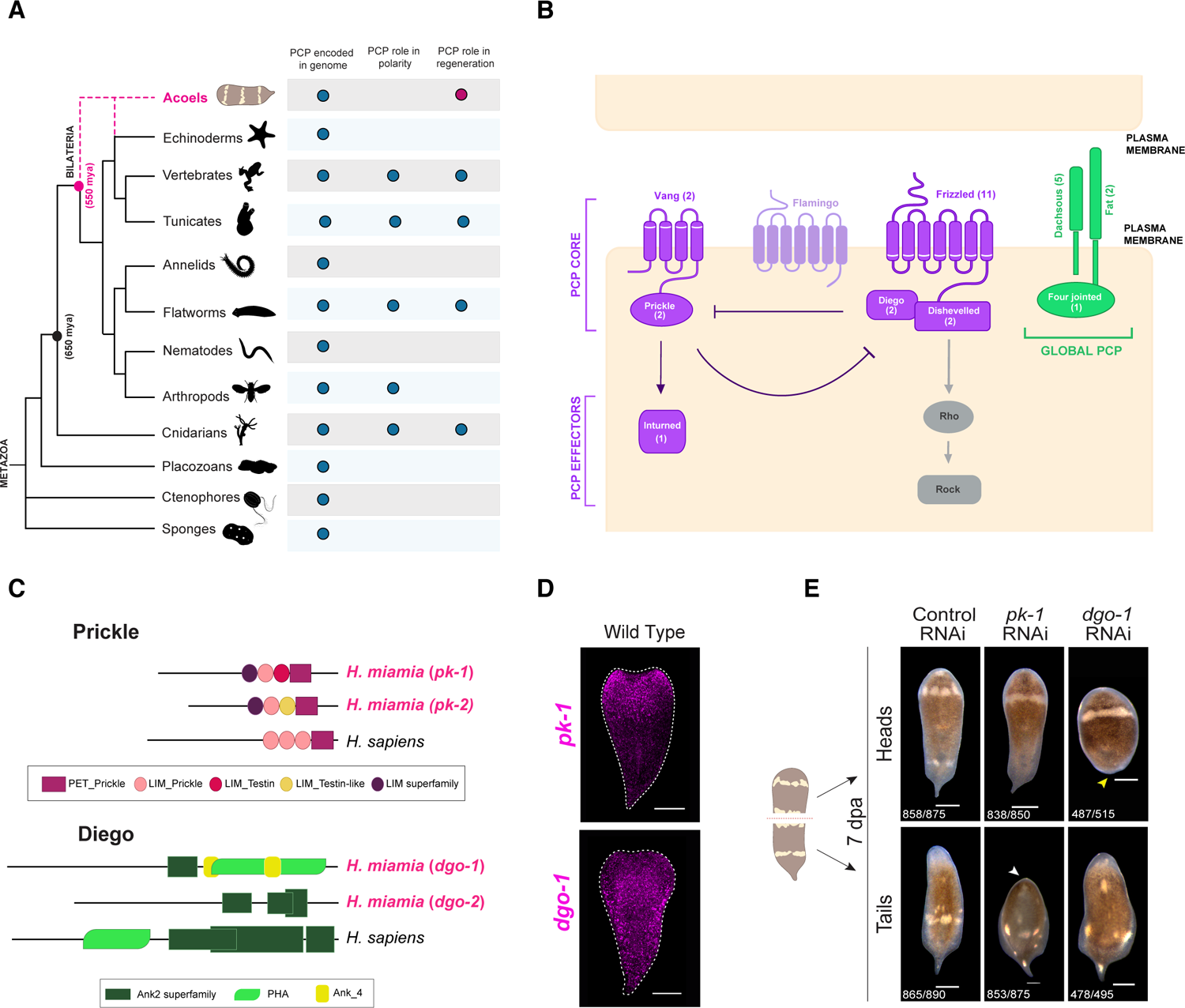
The genome of the acoel *Hofstenia miamia* encodes the majority of Planar Cell Polarity pathway components. (A) Schematic phylogenetic tree of major animal taxa highlighting the position of acoels as the sister group to all bilaterians diverging over ∼550 million years ago (mya) and from non-bilaterians over ∼650 mya. Dashed magenta lines indicate an alternative evolutionary scenario of acoels being closely related to echinoderms and hemichordates (Kapli and Telford, 2020; Philippe et al., 2011). Colored circles depict a specific trait present in at least one species of the indicated clade. The red circle denotes the findings of this study. All animal silhouettes from sponges to echinoderms were taken from PhyloPic. **(B)** Overview of the PCP signaling modules and interactions. The “core” complex (colored in purple) involves six members including three multipass transmembrane proteins: Flamingo, Vang and Frizzled, and three cytoplasmic proteins: Dishevelled, Diego and Prickle. A homolog for Flamingo was not found in *Hofstenia*. Global PCP signaling contains the large protocadherins Fat and Dachsous and the Golgi transmembrane kinase Four-jointed shown in green. PCP effector Inturned is specifically associated with the planar cell polarity whereas Rho and Rock effectors are not exclusive for PCP but are involved in numerous other cellular pathways. Numbers in parenthesis indicate the number of homologs found in *Hofstenia.* **(C)** Schematic representation of the protein domain composition for *pk-1* and *dgo-1* based on the closest human BLAST hit for the proteins. Black lines represent protein sequences and colored shapes indicate different protein families. **(D)** *In situ* hybridization expression patterns for *pk-1* and *dgo-1* in intact juvenile worms. **(E)** Asymmetric regeneration phenotypes from *pk-1* and *dgo-1* RNAi after transverse amputation. Control RNAi animals regenerated new heads and pointy tails by 7dpa. Both *pk-1* and *dgo-1* RNAi knockdown animals exhibited penetrant phenotypes resulting in wound closure but subsequent failure to regenerate heads (white arrowheads) and tails (yellow arrowheads) respectively. Conversely, *pk-1* anterior fragments regenerated new tails whereas *dgo-1* tail fragments produced new normal heads similar to control animals. Dashed red line in the schematic indicates the level of the transverse amputation. Scale bars: 100 µm.

Planar Cell Polarity (PCP) is a fundamental mechanism that collectively coordinates the directional organization of cells along the plane of a tissue. This phenomenon is predominantly regulated by two major sets of conserved proteins: the “core PCP’’ and “Fat/Dachsous (Ft/Ds)’’ pathways (Devenport, 2014; Goodrich and Strutt, 2011; Lapébie et al., 2011). PCP operates in a wide array of biological processes ranging from the alignment of hairs (trichomes), the arrangement of sensory bristles, and the configuration of the ommatidia of the compound eye in *Drosophila* (Jenny, 2010; Strutt, 2001) to convergent extension and limb bud elongation in vertebrates (Butler and Wallingford, 2017; Medina et al., 2000; Veeman et al., 2003; Yang and Mlodzik, 2015). PCP has also been implicated in regenerative processes in vertebrates; for example in the lateral-line system in zebrafish, the hair cell stereocilia bundles in the avian inner ear (Cooper et al., 2008; Dale et al., 2009; López-Schier and Hudspeth, 2006; Warchol and Montcouquiol, 2010), and during spinal cord regeneration in axolotls where it orients the axis of cell division along anterior-posterior axis (Rodrigo Albors et al., 2015) (Fig. 1A). Homologs of the Ft/Ds pathway components have polarized cellular localization in *Hydra* (Brooun et al., 2020) and homologs of the core PCP are needed for polarization of cilia in epidermal cells in planarians (Almuedo-Castillo et al., 2011; Vu et al., 2019). In the context of WBR, PCP components are involved in neural regeneration, where they limit neuronal growth (Beane et al., 2012). Although most components of core PCP and Ft/Ds pathways are present across metazoans, their roles in controlling cellular polarity and regeneration have not been investigated in many lineages including xenacoelomorphs (Fig. 1A).

We undertook a comprehensive and unbiased study of Wnt/PCP pathway components in *Hofstenia* by identifying putative homologs in the transcriptome and assessing their spatial expression patterns and functions in regeneration. We found two components, *prickle-1* (*pk-1*) and *diego-1* (*dgo-1*), to be required for head and tail regeneration, respectively. *pk-1* RNAi resulted in reduced expression levels of immediate wound response genes and of the anterior-specific wound-induced gene *fz-7* specifically in anterior-facing blastemas. *dgo-1* was needed for the wound-induced expression of the posterior gene *tf7l2*. Notably, we did not find clear evidence for a requirement for *pk-1* or *dgo-1* for expression of the earliest wound-induced anterior-posterior (AP)-asymmetric genes, *wnt-3* and *sp5*. Moreover, *pk-1* and *dgo-1* co-expression was highly enriched in a subset of muscle cells that also expressed *follistatin* (*fstl*), and *pk-1* RNAi resulted in a failure to regenerate correctly oriented and connected muscle fibers in the anterior-facing wound. Although we did not recover evidence for a role in controlling regeneration polarity, our data implicate Wnt/PCP components as important for wound-induced expression of some AP-specific genes, and for orienting muscle. Our work expands the study of PCP genes by revealing their functions in acoels, the sister-group to all other animals with bilateral symmetry, as well as their role in the process of whole-body regeneration and muscle organization.

## RESULTS

### The *Hofstenia* genome encodes the majority of Planar Cell Polarity pathway components

To study the components of the Wnt/PCP pathway, we identified homologs of the core and the Ft/Ds complexes in the *Hofstenia* genome (Fig. 1B). Some PCP components were present in single-copy, e.g. *inturned* (*intu*) (a downstream effector of the core pathway) and *four jointed (fjx1)*, whereas others were duplicated, e.g., we found two paralogs each for *vang*, *prickle* (*pk*), and *diego* (*dgo*). In the Ft/Ds pathway, we identified five *dachsous* cadherin-related genes (*ds*) and two putative *fat*-like homologs (*ft*) (Fig. 1B). Protein family composition analysis in these homologs revealed domain architectures that were largely similar to members of these protein families in other metazoans (Fig. 1 C; Fig. S1A; Supplementary Table 1). Moreover, 11 Frizzled (*fz*) receptors and two Dishevelled (*dvl*) genes had been previously reported in the *Hofstenia* genome (Srivastava et al., 2014). Although we were unable to identify clear homologs for *flamingo* (Fig. 1B; Fig. S1A), our analyses show that all the other major components of the core and Ft/Ds pathways are encoded in the *Hofstenia* genome, suggesting that these pathways could be functional in mediating cell polarity or other processes in xenacoelomorphs.

To further characterize the core PCP and Ft/Ds components, we determined their spatial expression patterns using fluorescent *in situ* hybridization (FISH) in intact juvenile worms. We recovered a range of expression patterns, from broad localization all over the body to differential expression along the anterior-posterior axis (Fig. 1D; Fig. S1B). For example, mRNAs for *dvl-1*, *vang-1*, and *pk-1* were enriched in the head or anterior region of the worm, whereas other genes such as *vang-2*, *ds-*like-1, *ds-*like-2, and *ds-*like-5 showed restricted expression in the posterior region or at the tip of the tail. Among genes that showed broadly distributed expression across the body (*dgo-1*, *dgo-2, intu*, *ds-*like-4, *fat-1*, *fat-2* and *fjx*), some exhibited prominent punctate expression patterns, e.g., *ds-*like-4, *fat-2* and *fjx1*. *ds-*like-3 showed a distinct mRNA distribution from those of other genes in this study, labeling cells at the edge of the body in the posterior and at the tip of the tail (Fig. S1B). The similarities in expression patterns suggested that these genes could work together in the same cells in *Hofstenia*, but the distinct expression patterns also suggest that these components may have independent roles as well.

Next, we sought to study the functions of these core PCP and Ft/Ds components. In *Hofstenia*, RNA interference (RNAi) applied during regeneration serves as a powerful tool to rapidly assess gene function as amputation forces the generation of new cells, which lack transcripts for the gene of interest. We injected worms with double-stranded RNA (dsRNA), amputated the body transversally, and screened head and tail fragments 7 days post amputation (dpa) to assess regeneration defects (Fig. 1E; Fig. S1C, D). 15 out of 17 genes assayed did not show a conspicuous effect or external abnormalities during regeneration upon RNAi, despite efficient knockdowns of target gene mRNA (Supplementary Table 2). Some RNAi head fragments for *vang-1* (n=4/16), *dgo-2* (n=7/30), and *intu* (n=6/16) seemed to have regenerated shorter tails but tail fragments produced new normal heads. Similarly, few tail fragments of *fjx1* RNAi (n=5/16) and *ds-*like-4 RNAi (n=3/16) appeared bloated but still were able to regenerate normal heads whereas *pk-2* RNAi worms seemed to have more elongated bodies (n=11/30) compared to control worms (Fig. S1C). Because these phenotypes were inconsistent across animals and the proportion of worms with the particular defect was low (not a penetrant phenotype), we categorized these genes as not having an impact on regeneration. Strikingly, loss of activity of *pk-1* and *dgo-1*, led to highly penetrant and distinct asymmetric phenotypes after RNAi treatment (Fig. 1E). Hence, we focused further analyses on these two genes.

### *pk-1* and *dgo-1* RNAi disrupt regeneration asymmetrically along the anterior-posterior axis

*pk-1* expression was highly enriched in the anterior compartment whereas *dgo-1* showed broad expression all over the body in intact animals (Fig. 1D), but both genes displayed dynamic expression during regeneration, with enrichment in head fragment wound sites at 24 and 48 hpa for *dgo-1* and *pk-1*, respectively, and in tail fragment wound sites starting at 15 hpa for both genes (Fig. S2A). This upregulation in both head and tail fragments contrasted with asymmetric regeneration outcomes – *pk-1* and *dgo-1* RNAi animals showed highly penetrant phenotypes with aberrant regeneration in anterior- and posterior-facing wounds, respectively. *pk-1* RNAi tail fragments were unable to form normal blastemas and subsequently failed to regenerate new heads after amputation (n= 853/875). However, loss of *pk-1* mRNA had no effect in head fragments, which restored new pointy tails similar to control worms (n= 838/850) (Fig. 1E).

Conversely, *dgo-1* RNAi head fragments failed to produce posterior tissues exhibiting no tail formation or incomplete tail regeneration (n=487/515). Although the *dgo-1* RNAi phenotype was robust, we observed that some animals formed unpigmented tissue outgrowth (blastema), sometimes with short pointy tails but did not make fully formed tails (Fig. 1E; Fig. S1D). In contrast, *dgo-1* RNAi tail fragments regenerated normal heads similar to control animals after 7 days of regeneration (n= 478/495) (Fig. 1E). Given that *pk-1* and *dgo-1* RNAi led to aberrant regeneration along the AP axis, we sought to characterize these defects using specific organ and axis region marker genes.

*dgo-1* RNAi fragments correctly expressed *fz-1* in posterior-facing wounds despite the absence of a clear tail (Fig. 2A). Since *dgo-1* RNAi heads were able to maintain tail identity but did not achieved posterior regeneration, we examined the expression of additional genes, *ds-* like-5, *fz-2*, *wnt-4* and *sp5*, expressed in the posterior in *Hofstenia* (Ramirez et al., 2020; Srivastava et al., 2014; Tewari et al., 2019). Surprisingly, we observed that *dgo-1* RNAi head fragments exhibited diminished expression levels for all the markers compared to control animals, albeit correctly specifying them in the appropriate location, suggesting the maintenance of correct posterior polarity in defective regenerated tails (Fig. 2B). Transverse amputations in our experiments removed the gut almost in its entirety from the anterior fragment, forcing the original head to re-establish its whole digestive system. *In situ* hybridizations for gut-specific markers, *aspp* and *pghd-4,* provided clear evidence of a reduced digestive cavity, consistent with these fragments’ gross failure to form new posterior structures (Fig. S2B). Using anterior molecular markers for differentiated tissues, we corroborated that there were no regeneration defects at the anatomical level in *dgo-1* RNAi tail fragments where animals were able to restore the anterior region of the body (Fig. S2C).

**Figure 2.**
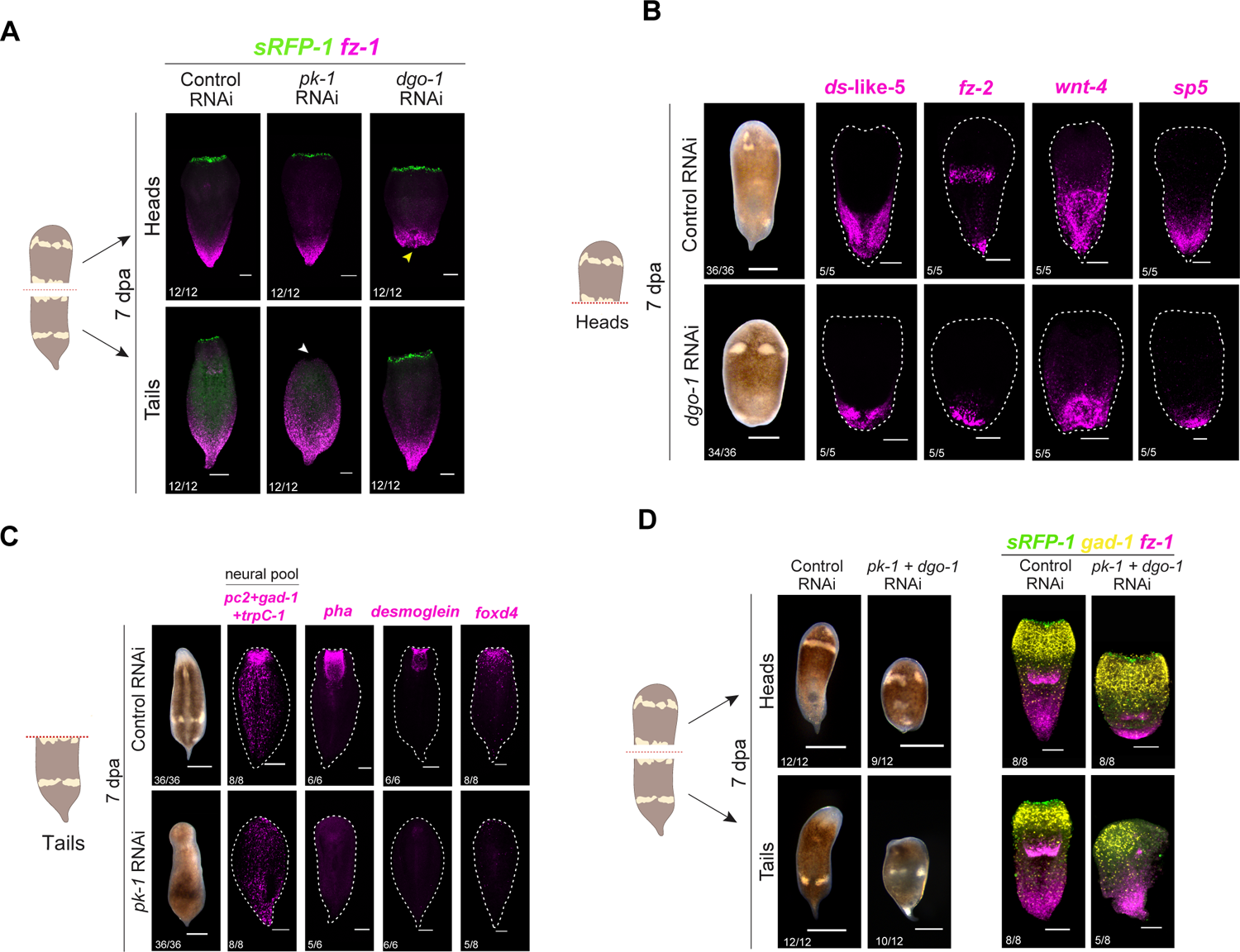
*pk-1* and *dgo-2* RNAi disrupt anterior-posterior regeneration in an asymmetric manner. **(A)** Double fluorescent *in situ* hybridization of anterior (*sFRP-1*) and posterior (*fz-1*) markers did not show a switch in polarity for *pk-1* and *dgo-1* RNAi animals. *pk-1* RNAi tails failed to produce anterior cell types expressing *sfrp-1* (white arrowheads) while *dgo-1* RNAi heads showed posterior expression of *fz-1,* despite no tail regeneration, indicating correct posterior specification (yellow arrowheads). **(B)** *dgo-1* RNAi head fragments displayed correctly polarized expression after wounding for multiple posterior marker genes (*ds*-like-5, *fz-2, wnt-4, sp5*) despite the lack of tail regeneration. **(C)** *pk-1* RNAi regenerating tail fragments lack expression of multiple neural anterior markers (*pc2, gad-1* and *trpC-1*) as well as mouth and pharyngeal marker (*desmoglein*, *pha*, *foxd4*), suggesting that *pk-1* is required to produce newly differentiated cells. **(D)** Simultaneous knockdown for *pk-1* and *dgo-1* in which head and tail double RNAi fragments reproduced the same phenotype as observed in the single knockdowns. Both fragments failed to regenerate new heads and tails and did not display any polarity defect or misexpression of anterior (*sFRP-1, gad-1*) and posterior (*fz-1*) markers assessed by triple *in situ* hybridization. All regenerating tail and head fragments are shown at 7dpa. Dashed red lines in the schematics indicate the level of the transverse amputation. Scale bars: 100 µm.

*pk-1* RNAi animals failed to re-establish *sFRP-1* expressing cells in the anterior-facing wounds indicating head regeneration defects, but head fragments making new tails looked indistinguishable from controls (Fig. 2A). Because *Hofstenia-sFRP-1* has a distinctive expression of few cells forming a ring around the entire mouth, the lack of *sFRP-1* could indicate anterior defects associated with the absence of a mouth; therefore, we assessed whether other anterior structures such as the brain and pharynx were affected. During *Hofstenia* regeneration, newly differentiated pharynx and neural cells can be detected approximately at 3 dpa (Hulett et al., 2020; Ricci and Srivastava, 2021; Srivastava et al., 2014). *pk-1* RNAi tail fragments failed to express pharynx markers (*desmoglein, pha*), and markers of various differentiated neural cell types (e.g., *gad-1, pc2, TrpC-1*), showing complete absence of anterior-specific structures and corroborating the role of *pk-1* in patterning the anterior compartment (Fig. 2C).

The observations that 1) *dgo-1* RNAi head fragments expressed posterior markers despite the absence of a tail and 2) no ectopic expression of AP markers was detected in any regenerating fragment of *pk-1* and *dgo-1* RNAi animals suggest that the failure to regenerate heads and tails was not caused by the inability to determine where heads and tails should be made. We further asked if the worms would be more susceptible to polarity defects if the function of *pk-1* and *dgo-1* was silenced simultaneously in these cell populations due to their asymmetric phenotype in individual knockdowns. Double knockdowns by injecting dsRNA against *pk-1* and *dgo-1* concurrently, resulted in similar head and tail defects with no evidence of loss of anterior-posterior identity. Head and tail fragments did not display more severe defects or any other effects compared to single knockdowns (Fig. 2D). Altogether, our results determined that *pk-1* and *dgo-1* activity is required to coordinate the correct AP patterning but axial polarity is not affected.

### *pk-1* RNAi animals show diminished expression of wound response genes during anterior but not posterior regeneration

To investigate the underlying mechanisms of the *pk-1* and *dgo-1* RNAi asymmetric phenotypes in an unbiased manner during *Hofstenia* regeneration, we applied bulk RNA sequencing (RNA-Seq) in head and tail fragments from control, *pk-1* and *dgo-1* RNAi animals. We focused this experiment on 40 hpa because 1) our prior bulk RNA-seq analyses showed that this time point captures the expression of wound response and patterning genes as well as of genes needed for stem cell differentiation, and 2) *pk-1* and *dgo-1* expression was upregulated at anterior and posterior facing wounds starting at 15 and 24 hpa, respectively (Fig. S2A). We reasoned that because *pk-1* and *dgo-1* are members of the core PCP pathway, they could be acting body-wide and their silencing would affect the whole worm; therefore, we aimed to evaluate the transcriptional changes occurring in the entire body by using complete head and tail fragments as opposed to isolating only the wound sites.

Differential expression analysis revealed significantly diminished expression of the wound-response program, activated immediately after injury, in *pk-1* RNAi tail fragments relative to controls (Fig. 3A; Fig. S3A). The wound response program, a gene regulatory network controlled by the early growth response factor Egr (Egr-GRN), is required for initiating regeneration and includes direct transcriptional targets of Egr such as *runt*, *follistatin* (*fstl*), *neuregulin* (*nrg-1*), and *metastasis suppressor-1* (*mtss-1*) also known as immediate early genes (IEGs). The Egr-GRN, was exclusively downregulated in *pk-1* RNAi tails, i.e., the fragments that were not able to regenerate new heads, but not in head fragments, which were able to regenerate normal tails (Supplementary Table 3A-B). This result is remarkable since the immediate wound-response program has been found to be transcriptionally upregulated in both head and tail fragments (Gehrke et al., 2019; Srivastava, 2021). Other wound induced genes, *traf-1* and *dus19,* also showed diminished expression in *pk-1* RNAi tail fragments. To corroborate these RNA-seq data, we quantified relative mRNA levels of genes affected under *pk-1* RNAi by RT-qPCR. First, we confirmed the efficiency of *pk-1* knockdown, finding significant reduction of the mRNA levels at 6 and 40 hpa (*p-value* < 0.001) compared to control fragments (Fig. 3B; Fig. S3B). Second, we found that the mRNA transcripts for *egr, runt*, *fstl*, *nrg-1*, and *mtss-1* were also significantly downregulated (*p-value* < 0.005) compared to control animals at 6hpa (Fig. 3B). Furthermore, we studied the expression of the Egr-GRN members in *pk-1* RNAi tail fragments via *in situ* hybridization. Control worms regenerating normal heads showed upregulation of these gene transcripts at the anterior-facing wounds at both time points while expression for the IEGs at the wound site was visibly diminished in *pk-1* RNAi animals at 6 hpa (Fig. 3C). At 40 hpa, we also detected diminished expression by *in situ* hybridization for the Egr-GRN, of which *egr* and *mtss-1* were significant reductions as validated by qPCR (Fig. 3C; Fig. S3B).

**Figure 3.**
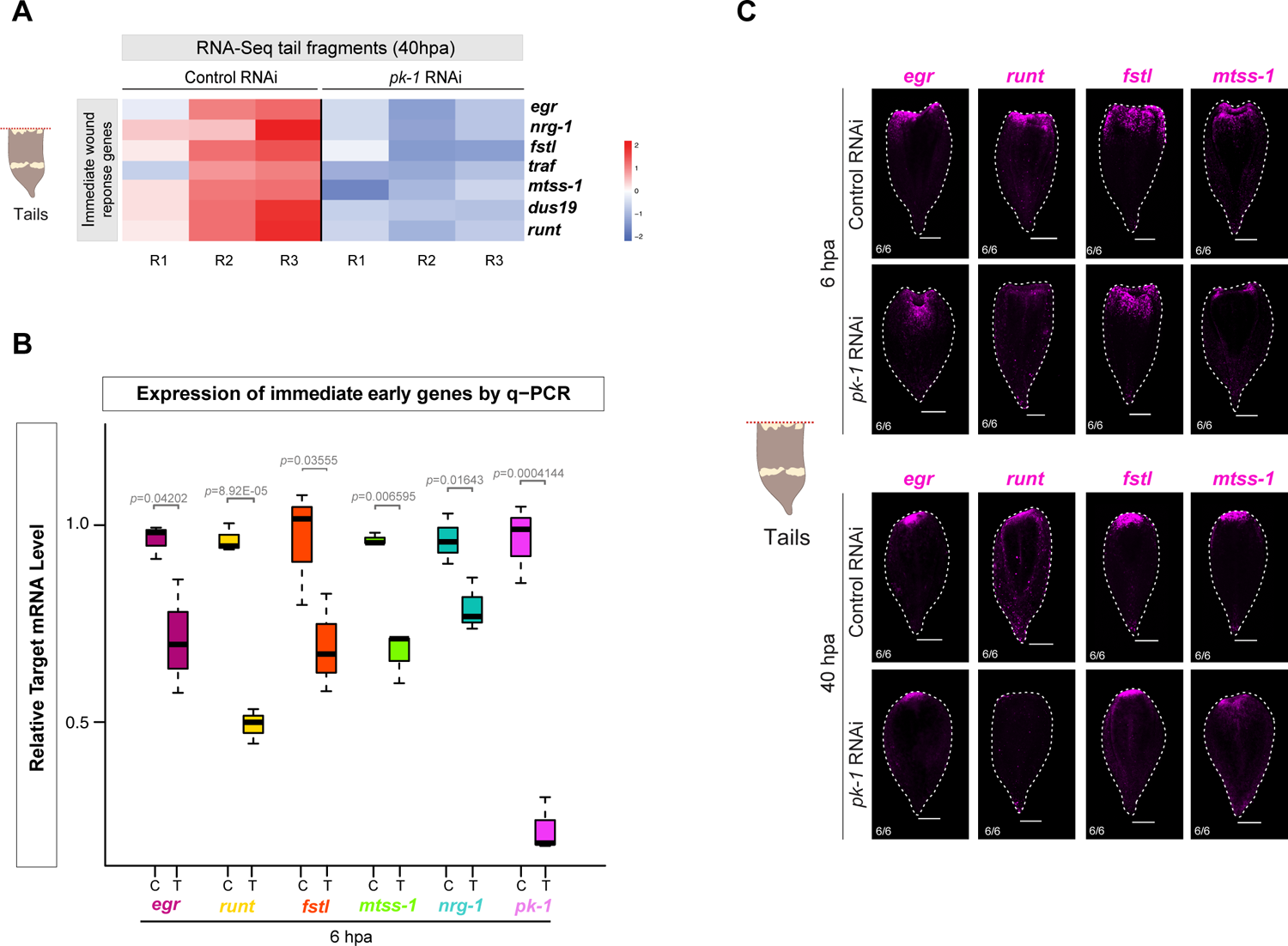
*pk-1* RNAi impacts the expression of wound response genes during anterior but not posterior regeneration. **(A)** Heatmap of RNA-Seq data comparing control vs *pk-1* RNAi tails at 40 hpa. Transcriptomic profiling of *pk-1* RNAi animals identified downregulation of genes associated with the immediate wound response. The RNA-Seq experiment included three biological replicates for control and treatment (R1, R2, R3) with three fragments per replicate. **(B)** qRT-PCR corroboration with primers specific to wound-induced genes *egr*, *runt*, *follistatin* (*fstl*), *metastasis suppressor* (*mtss-1*) and *neuregulin* (*nrg-1*) at 6 hpa, confirming the significant downregulation of the transcript levels. *pk-1* RNAi samples included three biological replicates per target gene with three fragments per replicate. Wilcoxon rank-sum test was used to calculate statistical significance (p-value <0.05). Housekeeping gene *GAPDH* was used for normalization. Control and treatment (RNAi) are indicated with C and T, respectively. **(C)** *In situ* hybridization for the immediate early genes downregulation in *pk-1* RNAi tails at 6 and 40 hpa. Dashed red lines in the schematic indicate the level of the transverse amputation. Scale bars: 100 µm.

### *pk-1* and *dgo-1* RNAi animals show diminished wound-induced expression of polarized Wnt pathway genes

In addition to revealing an impact on the Egr-GRN, our RNA-seq analysis for *pk-1 RNAi* tail fragments also identified effects on a cohort of genes associated with the Wnt signaling pathway which are known to be significantly upregulated at 12 hpa in anterior-facing wounds during *Hofstenia* regeneration (Ramirez et al., 2020) (Fig 4A, Fig. S3A; Supplementary Table 3). These targets correspond to sFRP genes (*sFRP-1*, *sFRP-2*), a Wnt ligand (*wnt-5*) and the anteriorly-expressed frizzled genes (*fz-7*, *fz-9*, *fz-11*). We experimentally corroborated via *in situ* hybridization that these genes failed to be expressed in *pk-1* RNAi (Fig. 4B). The lack of expression of this set of anterior-specific genes suggests that *pk-1* activity is necessary for reestablishing patterning information along the AP axis, consistent with failure to regenerate different tissues and organs in the head compartment. Given the impact on wound response genes in *pk-1* RNAi, we asked if the wound-induced expression of these patterning genes was also affected. Among this set of downregulated anterior expressed genes in *pk-1* RNAi, *fz-7* is the earliest to be expressed at the anterior-facing wound site (Ramirez et al., 2020). We found that *fz-7* was detected as wound-induced by 12 hpa in *pk-1* RNAi tail fragments, but at lower levels relative to control animals. This expression was detectable at 48h, again lower relative to controls, but was not sustained in 7 dpa in *pk-1* RNAi tail fragments, likely due to their failure to regenerate anterior tissues correctly (Fig. 4C). Notably, *wnt-3* and *sp5*, the earliest asymmetrically-expressed patterning genes, which can be detected in posterior but not anterior wound sites within 3 hpa, showed normal expression in *pk-1* RNAi.

**Figure 4.**
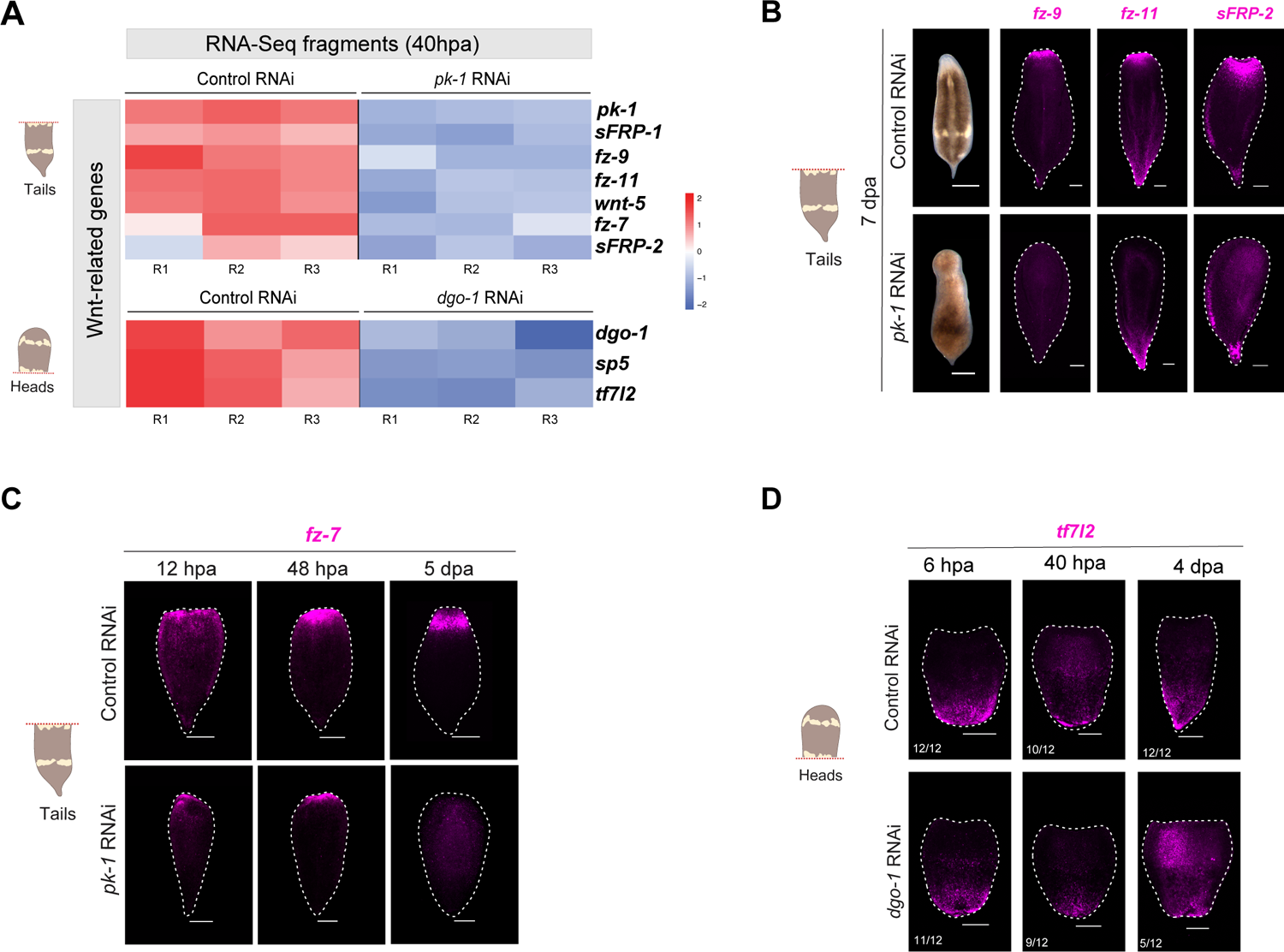
*pk-1* and *dgo-1* RNAi affect the wound-induced expression of polarized Wnt pathway genes during *Hofstenia* regeneration. (A) Heatmap of RNA-Seq data comparing control vs *pk-1* RNAi tails and control vs *dgo-1* RNAi heads at 40 hpa. Transcriptomic profiling of *pk-1* and *dgo-1* RNAi animals identified downregulation of Wnt-related genes known to be differentially induced at anterior-facing wounds between 6-12 hpa in tail fragments and posterior-facing wounds between 3-6 hpa in head fragments. *pk-1* and *dgo-1* were included in heatmaps to confirm the efficiency of the knockdowns by RNAi. The RNA-Seq experiment included three biological replicates for control and treatment (R1, R2, R3) with three fragments per replicate. (B) *In situ* hybridization confirms that Wnt-related genes failed to be re-expressed in the anterior part of the body corroborating the head regeneration defect and lack of anterior tissues in *pk-1* RNAi tails. (C) *In situ* hybridization of the anterior specific patterning gene *fz-7* at different timepoints of regeneration in *pk-1* RNAi animals. At 12 hpa, new expression of *fz-7* at anterior-facing wounds was observed in control animals but slightly diminished level of expression in *pk-1* RNAi tail fragments. *fz-7*^+^ cells were sustained to about 48 hpa but levels of expression were still much lower than control RNAi worms. Eventually, by 5 dpa, *fz-7* expression was completely lost, suggesting that *pk-1* does not have an early effect on turning on this specific anterior gene but plays a role in its maintenance for proper head regeneration. (D) *In situ* hybridization of the posterior specific patterning gene *tf7l2* at different timepoints of regeneration. Expression of *tf7l2* in posterior-facing wounds of *dgo-1* RNAi head fragments is diminished at 6 hpa but it is maintained at a later time points despite the inability of the worms to form new tails. Dashed red lines in the schematics indicate the level of the transverse amputation. Scale bars: 100 µm and 150 µm in panel (D).

Transcriptome profiling for *dgo-1* RNAi head fragments allowed us to identify individual differentially expressed genes, which did not include Egr-GRN members, contrasting with *pk-1* RNAi effects (Fig. S3C; Supplementary Table 3C-D). To understand the impact of *dgo-1* RNAi, we selected 24 significantly downregulated candidate genes following *dgo-1* RNAi in head fragments, based on conserved domains and moderate to high levels of transcript expression (>25 TPM in control RNAi heads), for further analysis (Supplementary Table 4). *In situ* hybridizations showed a variety of gene expression patterns, with many labeling neurons (*pc2*), putative sensory neurons (*zc21a*, *ci171*, *unk-240*, *wdr86*, *tenx-2*, *tm206-2*, *pkd2*), gut tissues (*pgdh4* and *plpl4*), cells in the anterior (*foxd4*, *sspo*, *unk-5327* and *gpcp1*), and cells in the posterior (*tf7l2*, *sp5*, *unk-587* and *lphn3*) (Fig. S3D). Head fragments of *dgo*-1 RNAi animals were able to restore the majority of the cell types, but the extent/pattern of expression differed from controls because of the failure of full tails to form (Fig. S3E); conversely, *dgo-1* RNAi tail fragments were able to fully regenerate anterior neural tissues (Fig. S3E). Based on these results, we hypothesized that *dgo-1* might regulate neural tissue turnover, rather than controlling neural regeneration. Our RNA-seq data analysis also showed a downregulation of *tf7l2* in *dgo-1* RNAi heads at 40 hpa (Fig. 4A; Fig. S3C). We focused on *tf7l2*, a posterior expressed gene belonging to the TCF/LEF family that are known mediators of Wnt signaling, which is robustly wound-induced by 6 hpa in posterior-facing wounds in *Hofstenia* (Gehrke et al., 2019; Ramirez et al., 2020). *In situ* hybridization showed diminished expression of *tf7l2* in posterior-facing wounds at 6 and 40 hpa in *dgo-1* RNAi head fragments, corroborating the RNA-seq data (Fig. 4D; Fig. S3C). *tf7l2* was not affected in *dgo-1* RNAi tail fragments regenerating new heads (Fig. S4A). Further, we found that *tf7l2* RNAi led to similar phenotypic outcomes as *dgo-1* RNAi (Fig. S4B), however, our results could not determine if *tf7l2* is a downstream target of *dgo-1* or if it is the *tf7l2* wound induce signal what is affecting tail regeneration. It is possible that the change in gene expression associated with *tf7l2* in *dgo-1* RNAi head fragments was a consequence of the inability to regenerate posterior tissues which is where *tf7l2* is typically expressed. Although *sp5*, a wound-induced gene whose expression precedes *tf7l12* in posterior wounds was affected in *dgo-1* RNAi in our RNA-seq study, we were still able to detect its expression by FISH (Fig. S4C). However, *wnt-3*, also wound-induced prior to *tf7l2*, was not detected as diminished in the *dgo-1* RNAi RNA-seq data or in the *in situ* hybridization experiments (Fig. S3C; Fig. S4C).

Altogether, the effects of *pk-1* and *dgo-1* RNAi on wound-induced Wnt pathway or axially restricted genes mirrored the effects on longer-term regeneration of tail and head fragments respectively. An anterior wound-induced gene, *fz-7*, showed diminished expression in tail fragments in *pk-1* RNAi but not in *dgo-1* RNAi (Fig. S4D), whereas a posterior wound-induced gene, *tf7l2*, was impacted in head fragments under *dgo-1* RNAi but not *pk-1* RNAi. Given that *pk-1* and *dgo-1* are upregulated at later time points of regeneration, their preexisting expression must be needed for expression of wound-induced Wnt pathway/AP patterning genes.

### *pk-1* is expressed in muscle and controls muscle fiber organization during regeneration

To understand how expression of *pk-1* and *dgo-1* prior to amputation could impact gene expression upon wounding, we examined the tissue types in which these genes were expressed. We first used the *Hofstenia* hatchling juvenile single-cell RNA-seq atlas (Hulett et al., 2023) to determine the putative tissue-specific expression of all the PCP genes identified here. Remarkably, although the PCP is known to operate canonically in epidermis, our results indicated that only 24% of the PCP homologs were robustly expressed in the epidermis in *Hofstenia*. The majority of genes (65%), including *pk-1* and *dgo-1*, were found to be highly enriched in muscle (Fig 5A). *pk-1* was also expressed in neural and digestive cells types whereas *dgo-1* was additionally expressed in epidermal cells (Fig. 5A; Fig.S5A). Notably, in our *in situ* hybridization studies of *pk-1* expression, this gene exhibited stronger signal in the head compartment dorsally and around the mouth ventrally, which may be associated with neural tissues (Fig.1C; Fig.S5A). In contrast, *dgo-1* showed a broadly distributed expression pattern dorsally, with only a few *dgo-1*+ cells displaying a posterior gradient when viewed ventrally. Visualization of deeper tissue layers revealed stronger subepidermal signal throughout the body rim, possibly representing muscle cells (Fig.S5A).

**Figure 5.**
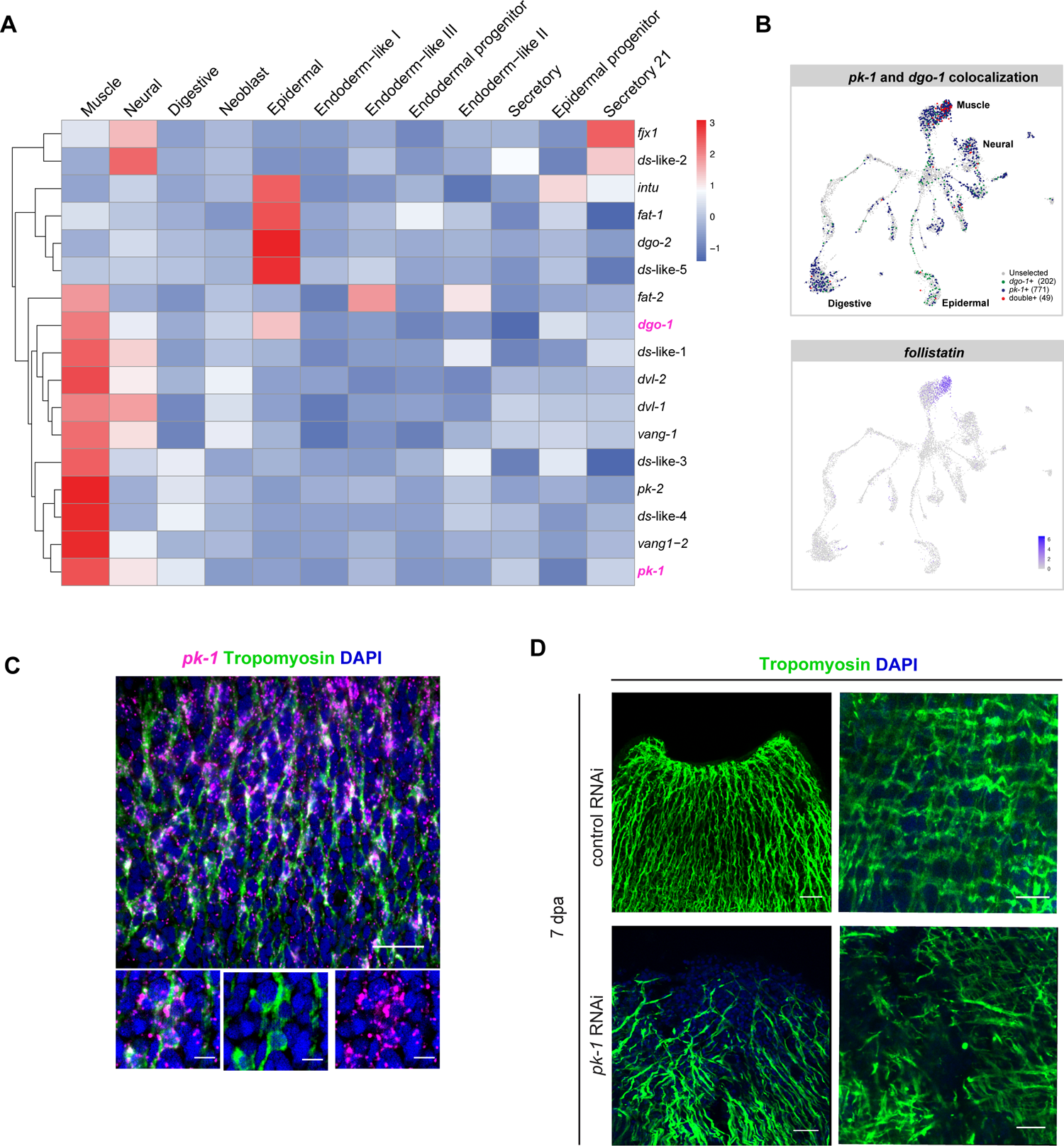
*pk-1* is expressed in muscle and causes muscle fiber disorganization defects during regeneration. **(A)** Heat map of cell type–specific clusters from single-cell RNA-seq atlas (Hatchling juvenile dataset) showing expression profiles of the 17 homologs of the Planar Cell Polarity pathway (PCP). The majority of the genes are expressed in muscle and neural tissues followed by expression in the epidermis. **(B)** Projection of *pk-1* and *dgo-1* into the single-cell RNA-seq atlas showing gene expression of each gene in different cell-type cluster and indicating colocalization for both genes specifically in the muscle cluster (top). Projection of *follistatin* into the single-cell RNA-seq atlas showing gene expression enrichment particularly in the muscle cluster where *pk-1* and *dgo-1* colocalize (bottom). **(C)** *In situ* hybridization for *pk-1* coupled with immunostaining employing *Hofstenia* custom Tropomyosin antibody displaying colocalization in the longitudinal muscle. **(D)** *pk-1* RNAi caused severe defects in muscle fiber regeneration and organization in the anterior region showing misaligned longitudinal muscle fibers and disorganization of the body wall muscle network with a lack of fibers at the anterior tip of the fragment. Scale bars 20 µm in (C), 25 µm in (D) left top and bottom panels and 15 µm in (D) right top and bottom panels.

Prickle and Diego are both cytoplasmic proteins required for the regulation of PCP activity, are asymmetrically localized in opposite sides of the cell and compete with one another for Dishevelled binding (Jenny, 2003; Jenny et al., 2005; Strutt and Strutt, 2009). To identify cells in which *pk-1* and *dgo-1* were co-expressed, we projected their expression onto the hatchling juvenile single-cell RNA-seq atlas. Of 771 *pk-1*^+^ cells and 202 *dgo-1*^+^ cells total, 49 *pk-1*^+^ (6.7%) and *dgo-1*^+^ (24%) cells showed co-expression in different cell-type clusters with 31 of the 49 *pk-1*^+^/*dgo-1*^+^ cells found in the differentiated muscle cluster (Fig. 5B). In particular, these *pk-1*^+^/*dgo-1*^+^ cells were enriched in a subset of muscle cells that are marked by the expression of *fstl*, which is expressed in longitudinal muscle (Gehrke et al., 2019) (Fig. 5B). Notably, in addition to observing the impact on wound-induced expression of *fstl* in *pk-1* RNAi tail fragments (Fig. 3A-C), we found that *fstl* expression was absent at the anterior-facing wound at 7 dpa, suggesting that *pk-1* is needed for regeneration of *fstl*^+^ muscle (Fig. S5B). *fstl* was not affected under *dgo-1* RNAi.

To corroborate the enrichment of *pk-1* and *dgo-1* within muscle tissue, we implemented *in situ* hybridization to detect *pk-1* transcripts coupled with immunostaining employing a *Hofstenia* custom Tropomyosin antibody. While we were unable to resolve cellular level expression of *dgo-1*, we found robust co-expression of *pk-1* with tropomyosin confirming that *pk-1* is expressed in muscle fibers (Fig. 5C; Fig. S5C). This localization is consistent with our finding that *pk-1* RNAi impacts Egr-GRN members and anterior Wnt pathway genes, many of which are expressed in muscle (Hulett et al., 2023b; Ramirez et al., 2020; Raz et al., 2017).

Therefore, we investigated muscle tissue morphology in *pk-1* RNAi fragments using the Tropomyosin antibody and found a substantial disorganization of the outer layer of longitudinal fibers and body wall muscle in the anterior of *pk-1* RNAi tail fragments whereas muscle regeneration in the new tails of *pk-1* RNAi heads was normal (Fig. 5D; Fig. S5D,E). Although *dgo-1 RNAi* heads were unable to regenerate new tails, the muscle at the wound area was not missing or disorganized as seen in *pk-1* muscle defect (Fig. S5E). Based on these data, we hypothesize that *pk-1* is needed for muscle organization, which is then needed for correct expression of wound induced genes including the Egr-GRN and Wnt pathway genes.

## DISCUSSION

We sought to study whether non-canonical Wnt signaling, in the form of Wnt/PCP signaling, and planar cell polarity pathways more generally, including both core PCP and Ft/Ds, control regeneration polarity in the AP axis in the acoel *Hofstenia miamia*. We did not find evidence for this, as animals with aberrant regeneration in functional studies did not display misspecification of anterior or posterior tissues. However, we found polarized regeneration outcomes, where *pk-1* was needed for anterior/head regeneration and *dgo-1* was needed for posterior regeneration, implicating these core PCP pathway components as having important roles in whole-body regeneration. These regeneration phenotypes potentially originate from problems with early stages of regeneration – wound response genes and the wound-induced anterior marker *fz-7* showed diminished expression in tail fragments under *pk-1* RNAi, and the wound-induced posterior marker *tf7l2* was diminished in head fragments under *dgo-1* RNAi. While we are unable to explain why these genes only impact regeneration of heads and tails, respectively, we found that *pk-1* and *dgo-1* are expressed in many cell types, but only co-expressed in longitudinal muscle. Furthermore, *pk-1* RNAi resulted in disorganized muscle specifically in the anterior wound site.

The impact of *pk-1* RNAi on muscle organization in *Hofstenia* is reminiscent of the impact that abrogation of non-canonical Wnt signaling has on muscle during regeneration in planarians (Gittin and Petersen, 2022). Wnt11 and Dishevelled, acting via non-canonical Wnt signaling, control the orientation of longitudinal muscle and the polarized wound-induced expression of *notum* in the planarian *Schmidtea mediterranea*. Given that Wnt ligands and Dishevelled can act in multiple Wnt-associated pathways, our data raise the possibility that Wnt/PCP could be the relevant non-canonical pathway for controlling regeneration polarity in planarians. However, contrasting with the planarian data, we did not find evidence for an effect of *pk-1* or *dgo-1* RNAi on regeneration polarity, as wound-induced indicators of polarity such as *wnt-3*, *sp5*, *tf7l2*, and *fz-7* were not expressed ectopically in head or tail wound sites. Instead, we found that some of these markers failed to be expressed in their expected wound sites of expression. This could suggest that, in acoels, Wnt/PCP components are needed for certain aspects of early anterior-posterior patterning, but do not play a role in the upstream polarity decision-making as other factors whose knockdown leads to a polarity reversal. Interestingly, knockdown of *ddx24* in planarians, which results in disorganized muscle, also impacts Wnt gene expression (Sarkar et al., 2021). While the relationship of this gene to Wnt11 and Dishevelled, or to PCP components, remains to be determined, the planarian and acoel data together point to a central role for well-oriented muscle in mediating regeneration of correctly polarized and patterned animals.

The anterior- and posterior-regeneration specific phenotypes of *pk-1* and *dgo-1* RNAi animals remain unexplained. It is possible that the differential phenotypes we observed are the result of different strengths of RNAi in the two perturbations, or that these genes do not operate in the same cells to control cell polarity. Studies of intracellular protein localization are needed to assess if Prickle and Diego proteins localize to opposite cell membranes of the same cell and if they control cellular polarity in *Hofstenia*. Interestingly, whereas PCP components have been shown to control epithelial cell polarity in planarians, the roles of these genes in early stages of regeneration are not known (Almuedo-Castillo et al., 2011; Vu et al., 2019).

Strikingly, *pk-1* and *dgo-1* were co-expressed predominantly in one cell type in *Hofstenia* – *fstl*^+^ muscle, which is a subset of longitudinal muscle. This, together with the disorganized muscle observed in *pk-1* RNAi animals, raises the possibility that core PCP components may be involved in orienting/polarizing muscle in acoels. Planar cell polarity operates in many contexts in epithelia, and has been shown to orient mesodermal/mesenchymal cells in vertebrates, but a role in orienting muscle has not yet been reported. Future work dissecting the mechanisms of PCP components in muscle in *Hofstenia* and other systems will enable us to assess the prevalence of polarized muscle across metazoans.

## METHODS

### Animal culture

*Hofstenia miamia* is a marine worm cultured under laboratory conditions in artificially synthesized sea water (ASW). Water was prepared at 37ppt with a pH in the range of 7.9 - 8. using Instant Ocean Sea Salt. Adult worms were kept in plastic boxes at 21°C containing approximately 1L of filtered ASW where they reproduced and laid embryos on the surfaces of the boxes. Embryos were collected using a glass pasteur pipette and raised in petri dishes for about 8 days at 24°C. Hatchling and juvenile worms were raised in zebrafish tanks at room temperature and fed with rotifers *Brachionus plicatilis*. Twice a week, adult *Hofstenia* boxes were cleaned, seawater was replaced, and worms were fed with freshly hatched brine shrimp *Artemia* sp. Animals used for experiments were starved for 3-5 days prior to regeneration assays.

### RNA interference (RNAi) Experiments

Double-stranded RNA (dsRNA) was synthesized following a standard protocol previously established in the lab and described in previous work (Gehrke et al., 2019; Hulett et al., 2023b; Srivastava et al., 2014). RNAi injection experiments were performed by introducing dsRNA corresponding to the target gene into the tail area of the worm for 3 consecutive days. dsRNA injections were made using a Drummond Nanoject II. Animals were cut transversally at least 2 hours after the last injection on the third day and fragments were allowed to regenerate for the appropriate number of days while being monitored for visible phenotypes or external defects. Control dsRNA for gene inhibition was the *unc22* sequence from *C. elegans* that is absent in *Hofstenia*. Phenotype screening and images were made using a Leica DM8000 stereoscope.

### Fixation and Fluorescent *in situ* Hybridization (FISH)

Whole worms and regenerating fragments at different timepoints upon amputation were fixed in 4% paraformaldehyde in 1% phosphate buffered saline with 0.1% Triton-X (PBST) for one hour at RT on a nutator. Three washes of 5 min each were made with PBST and animals were moved to a 1:1 mixture of PBST:methanol to gradually start dehydrating the tissue (5min).

Animals were transferred into a final change of 100% MeOH and stored at −20°C until use. Digoxigenin and Fluorescein labeled RNA probes for detection of mRNA expression were synthesized as previously described (Srivastava et al., 2014). Fluorescence *in situ* hybridizations (FISH) were performed following a detailed protocol established for *Hofstenia* (Srivastava et al., 2014) followed by minor modifications (Hulett et al., 2023b; Ramirez et al., 2020). Specimens were mounted in Vectashield and whole mounts were analyzed and imaged with a Leica SP8 confocal microscope. Images were processed in FIJI.

### Quantitative real time RT-PCR (qPCR)

RNA was extracted using the Nucleospin RNA XS kit (740902.10, Macherey-Nagel # 740902). Extracted RNA was used to synthesize cDNA by reverse transcription using the SuperScript III kit (Thermo Fisher 18080044) with random hexamer and oligoDT priming. Expression of target genes was detected with the SsoFast EvaGreen Supermix (Bio-Rad #172-5203) and the BioRad CFX96 Real-Time PCR System. Amplifications with forward and reverse primers were averaged across replicates and normalized by expression levels of a housekeeping gene (*gapDH*) to obtain a ΔCt value. Gene specific primers were designed using Primer3. A Wilcoxon rank sum test was used to determine the significance of mRNA level change between control and experimental RNAi conditions. We used the standardized threshold of *p* < 0.05 to determine statistical significance.

### RNA-sequencing (RNA-seq) Analysis

RNA-seq experiments were performed as biological replicates in triplicate by pooling 3 regenerating fragments per condition per replicate. Total RNA was extracted using the NucleoSpin RNA XS kit (Macherey-Nagel # 740902, without the use of carrier RNA), and the Illumina TruSeq Kit (Illumina #RS-122-2001) was used to generate sequencing libraries. Single-end 40- or 75-bp reads were obtained via Illumina HiSeq 2000 or NextSeq. Demultiplexed reads were quasi-mapped to the transcriptome using Salmon (Patro et al., 2017). Differential expression analysis was conducted separately for head and tail fragments and pairwise comparisons were made using sleuth and DeSeq2 between control RNAi and treatment RNAi (Pimentel et al., 2017; Varet et al., 2016) with default parameters. Genes with *p-*values <0.05 in both the posterior and anterior datasets were considered to be significantly differentially expressed. Single cell analysis and gene projection was performed using Seurat v.4 (Hao et al., 2021; Hulett et al., 2023).

---

**Supplementary Figure 1.** Characterization of the PCP pathway components in *Hofstenia*. **(A)** Domain composition analysis of Planar Cell Polarity (PCP) Pathway proteins found in *Hofstenia miamia* showing a conserved architecture across different invertebrate and vertebrate organisms (protein sequences from model organisms obtained from GenBank. Accession numbers provided in Supplementary Table 1). *Hofstenia* homologs in pink. **(B)** Gene expression patterns of PCP members found in *Hofstenia* detected by fluorescent *in situ* hybridization in juvenile worms. mRNA localization was observed in different domains along the anterior-posterior axis and scattered through the body. **(C)** Functional analysis of PCP genes after RNAi. Head and tail phenotypes were screened at 7dpa (days post amputation). Some animals showed defects but they were low in numbers and not consistent across experiments. *vang-1* (n=4/16), *dgo-2* (n=7/30) and *intu* (n=6/16) seemed to have shorter tails (yellow arrowheads) compared to control animals. Tail fragments for *ds-*like-4 (n=3/16) and *fjx1* (n=5/16) appeared bloated (white arrowheads) whereas *pk-2* (n=11/30) showed shorter head blastemas and elongated bodies (red arrowheads). Excluding these cases, animals for most of the genes did not produce visible external phenotypes, regenerating normal heads and tails. **(D)** Range of phenotypes observed in *dgo-1* RNAi head fragments displayed indented blastemas, fluctuating from small outgrowths of colorless tissue (white arrowheads) to truncated pointy tails (yellow arrowheads), but did not fully pattern the posterior portion of the body. Dashed red lines in the schematics indicate the level of the transverse amputations. Regenerating head and tail fragments are shown 7dpa. Scale bars: 100 µm.

**Supplementary Figure 2. *pk-*1 and *dgo-1* display dynamic expression during regeneration. (A)** Regeneration time course in wild type worms for *pk-1* and *dgo-1* revealed wound-induced expression for both genes in anterior-facing wounds at 15 hpa (white arrowheads). Posterior-facing wounds showed expression at 24 hpa for *dgo-1* (yellow arrowheads) whereas *pk-1* was upregulated at 48 hpa around the new regenerated tail (red arrowheads). **(B)** Gut markers in *dgo-1* RNAi showed failure to form new digestive system. **C)** Specific anterior markers for neural, mouth and pharynx tissues showed appropriate expression in the anterior compartment in *dgo-1* RNAi tail fragments indicating normal head regeneration. Regenerating tail fragments in (B) and (C) are shown at 7dpa. Dashed red lines in the schematics indicate the level of the transverse amputations. Scale bars: 100 µm.

**Supplementary Figure 3. *pk-1 and dgo-1* display distinct phenotypes during *Hofstenia* regeneration. (A)** Heatmap of RNA-Seq analysis for *pk-1* RNAi tail fragments at 40 hpa, showing statistically significant downregulation in gene expression for several genes including the early wound response program (highlighted in yellow) and polarity Wnt-related genes (highlighted in green) that are normally expressed at anterior-facing wounds in wild-type regenerating animals. The RNA-Seq experiment included three biological replicates for control and treatment (R1, R2, R3) with three fragments per replicate. **(B)** qRT-PCR corroboration with primers specific to wound-induced genes *egr*, *runt*, *follistatin* (*fstl*), *metastasis suppressor* (*mtss-1*) and *neuregulin* (*nrg-1*) at 40 hpa. *egr* and *mtss-1* showed significant reductions at this time point. *pk-1* RNAi samples included three biological replicates per target gene with three fragments per replicate. Wilcoxon rank-sum test was used to calculate statistical significance (*p*-value <0.05). Housekeeping gene *GAPDH* was used for normalization. Control and treatment RNAi are indicated with C and T, respectively. **(C)** Heatmap of RNA-Seq analysis for *dgo-1* RNAi head fragments at 40 hpa, showing statistically significant downregulation in gene expression for the wound-induced genes *tf7l2* and *sp5* (highlighted in yellow) and several neural-related genes. The RNA-Seq experiment included three biological replicates for control and treatment (R1, R2, R3) with three fragments per replicate. **(D)** *In situ* hybridizations in intact worms of *dgo-1* RNAi target genes selected from the head fragments RNA-Seq differential expression analysis based on the levels of downregulation compared to control RNAi (>25 transcripts per million). These genes are expressed in spatial patterns consistent with expression in putative sensory neurons, gut tissues, and in cells specific to the anterior or the posterior. **(E)** Validation of some of *dgo-1* RNAi target genes by *in situ* hybridization showing differences in posterior expression compared to control animals due to the lack of tail formation in *dgo-1 RNAi* (top). *dgo-1* RNAi tail fragments were able to regenerate normal heads expressing anterior neural cell types (bottom). Dashed red lines in the schematics indicate the level of the transverse amputations. Scale bars: 100 µm.

**Supplementary Figure 4.** *dgo-1* RNAi impacts the re-establishment of posterior tissue patterning during *Hofstenia* regeneration. (A) Left: Fluorescent *in situ* hybridization of the posterior specific gene *tf7l2* at different timepoints of regeneration in *dgo-1* RNAi tail fragments. *tf7l2* showed some expression at the anterior-facing wound in all animals indicating no differences between control and RNAi treatment. Regenerating tail fragments correspond to head fragments in Fig. 4D. Right: Chromogenic *in situ* hybridization was implemented to evaluate early *tf7l2* expression in *dgo-1* RNAi head and tail fragments specifically at 6hpa. The expression of *tf7l2* is clearly reduced at posterior-facing wounds in *dgo-1* RNAi heads compared to control animals corroborating the RNA-Seq and FISH results. No difference in *tf7l2* expression was observed between control and RNAi treatment in tail fragments. (B) *tf7l2* RNAi phenocopied the posterior defect observed in *dgo-1* RNAi after transverse amputation. Control RNAi animals regenerated new heads and pointy tails whereas *tf7l2* RNAi animals failed to regenerate new tails (white arrowheads) but produced new normal heads similar to control animals (yellow arrowheads) by 7dpa. *tf7l2* RNAi head fragments displayed phenotypes that range from no blastema formation to small outgrowths of colorless tissue or truncated pointy tails; however, these head fragments did not fully pattern the posterior portion of the body. (C) *In situ* hybridization for the posterior patterning genes *sp5* and *wnt-3* in control, *pk-1* and *dgo-1* RNAi regenerating head fragments at 0, 3 and 15 hpa. Wound-induced expression for these genes is observed at 3hpa and maintained at a later time points indicating that early patterning information is activated correctly in *dgo-1* RNAi heads with defects in tail regeneration. *pk-1* RNAi heads were included as an additional control since they regenerated proper tails. (D) *In situ* hybridization of the anterior specific patterning gene *fz-7* in *dgo-1* RNAi tail fragments showing correct specification in the new regenerated head. Dashed red lines in the schematics indicate the level of the transverse amputation. Scale bars: 100 µm and 150 µm in (C).

**Supplementary Figure 5. *pk-1* is required for correct muscle fiber organization during *Hofstenia* regeneration. (A)** Projection of *pk-1* and *dgo-1* into the hatchling juvenile single-cell RNA-seq atlas indicating their expression in different cell type tissues. *pk-1* is expressed in muscle, neural and digestive tissues whereas *dgo-1* is found in muscle and epidermis. **(B)** *In situ* hybridization for *fstl* in *pk-1* RNAi tails from dorsal and ventral views, showing the lack of expression of this gene in the anterior part of the body after 7dpa. **(C)** *In situ* hybridization for *pk-1* coupled with immunostaining employing *Hofstenia* custom Tropomyosin antibody displaying colocalization mainly in the longitudinal muscle. This experiment corresponds to Fig. 5C. **(D)** *pk-1* RNAi caused severe defects in muscle fiber regeneration including the lack of a mouth and associated musculature. **(E)** General view of muscle architecture along the AP axis in control, *pk-1* and *dgo-1* RNAi head and tail fragments. Scale bars 100 µm in (A) and (B), 20 µm in (C), 25 µm in (D) and 200 µm in (E).

## ACKNOWLEDGEMENTS

We thank all the former members of the Srivastava and Koenig Labs for their continued support and valuable discussions during the initial phase of this study. Thank you to all the current members of the Srivastava Lab for their constant encouragement, helpful feedback and critical input. Special thanks to Dr. Andrew R. Gehrke for assistance provided with the RNA-Seq data analysis and Drs. Allison Kann and Ye Duan for their guidance on protein domain predictions. We extend our gratitude to Dr. Paul Bump and Catriona Breen for providing valuable comments and suggestions for this version of the manuscript. D.M.B. was supported by AAUW American Postdoctoral Fellowship and National Science Foundation (award 1652104). N.R.-R. was supported by a ‘Formación del Profesorado Universitario (FPU13/01840)’ PhD fellowship from the Spanish ‘Ministerio de Educación, Cultura y Deporte (MECD)’, a Boehringer Ingelheim Fonds travel grant, and a travel scholarship from the Spanish Society of Genetics. M.S. was supported by the Searle Scholars Program, Smith Family Foundation, and National Science Foundation (award 1652104).

## Notes

### Competing Interest Statement

The authors have declared no competing interest.

