## Supplementary figures and images for "PCP components control anterior and posterior regeneration, with a Prickle homolog impacting muscle organization, in the acoel *Hofstenia miamia*"

### Bolanos_etal_SupplFig_bRxV_100323.pdf

#### Supp. Figure 1

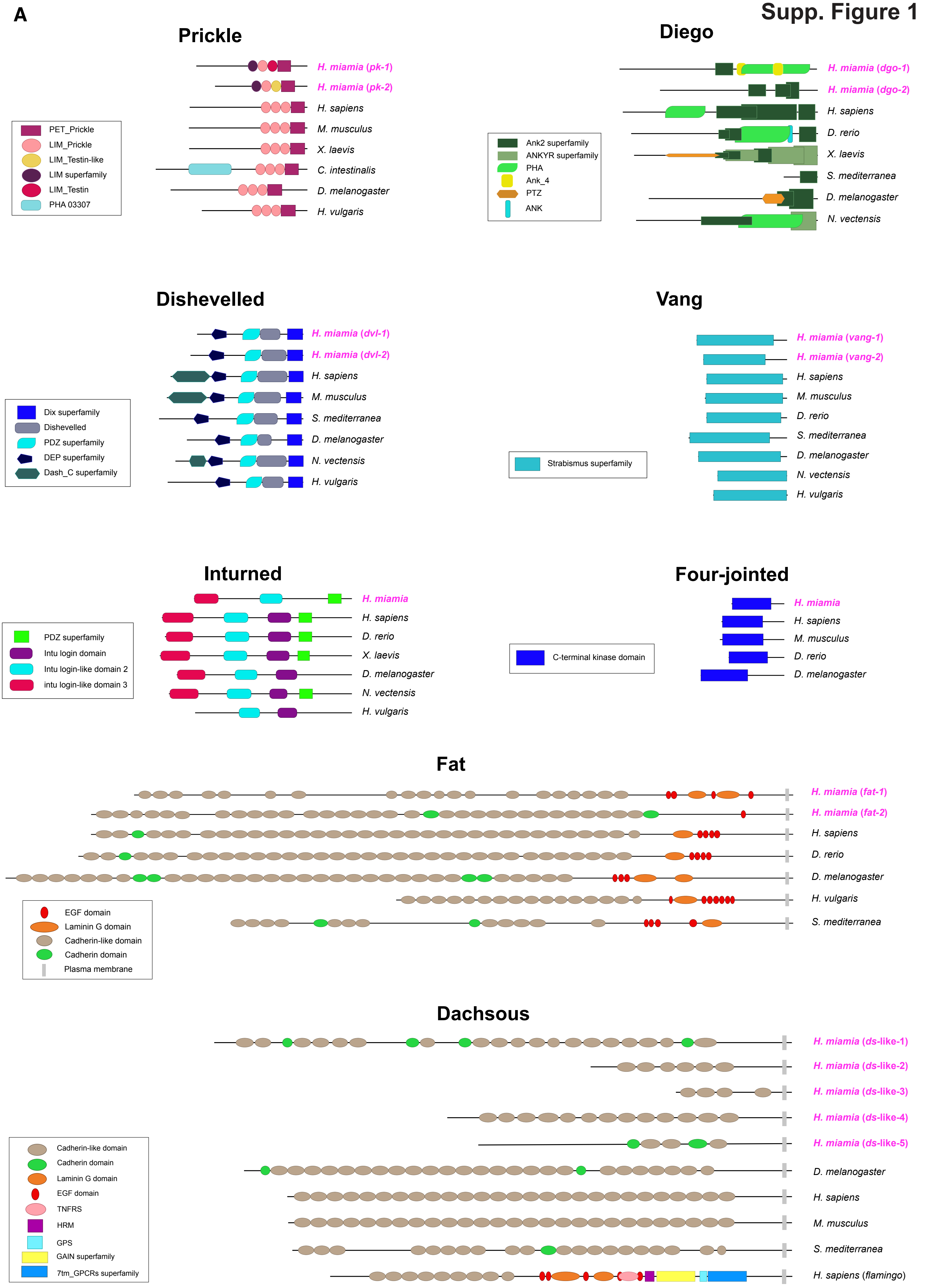

Supp. Figure 1

B

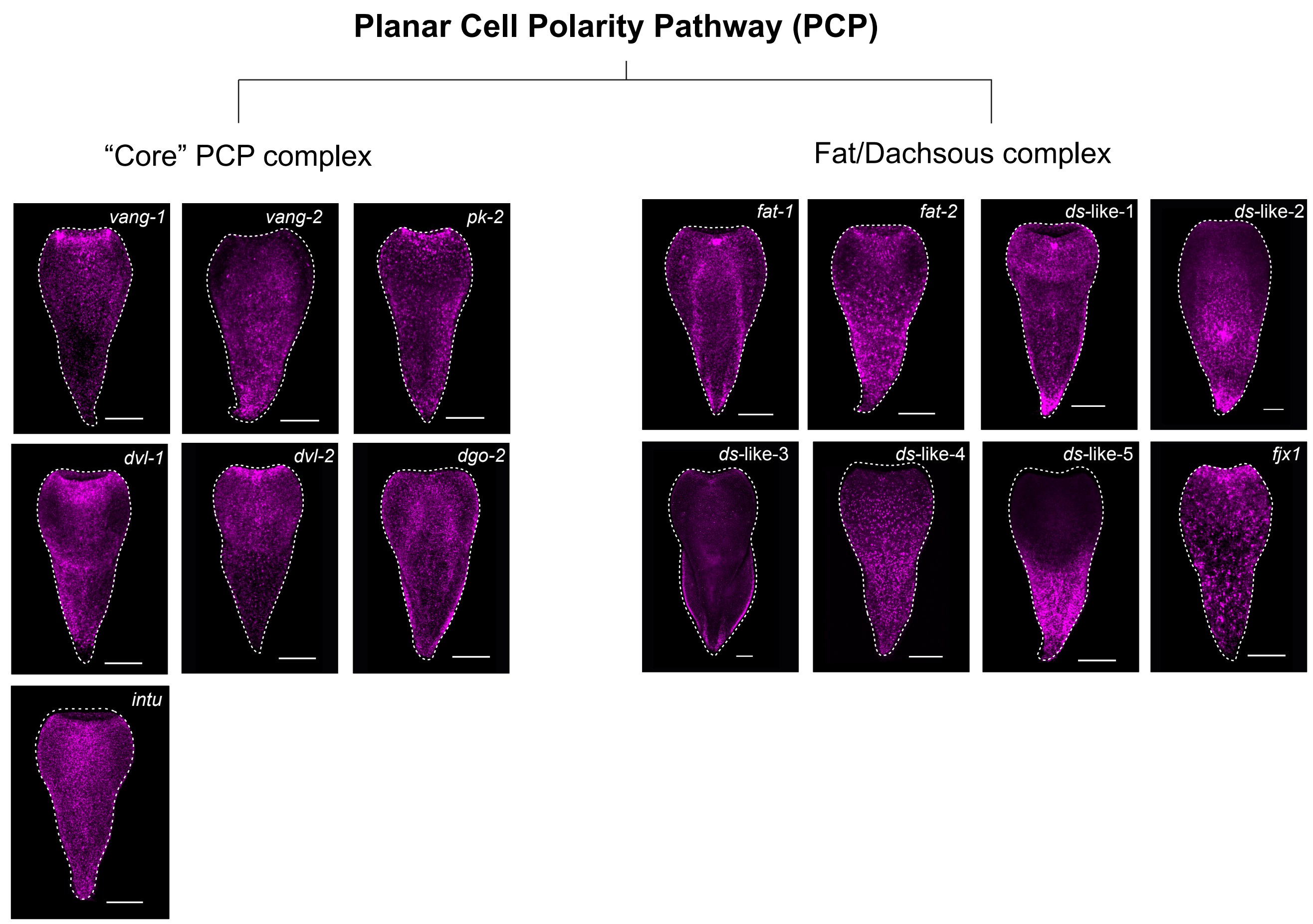


C

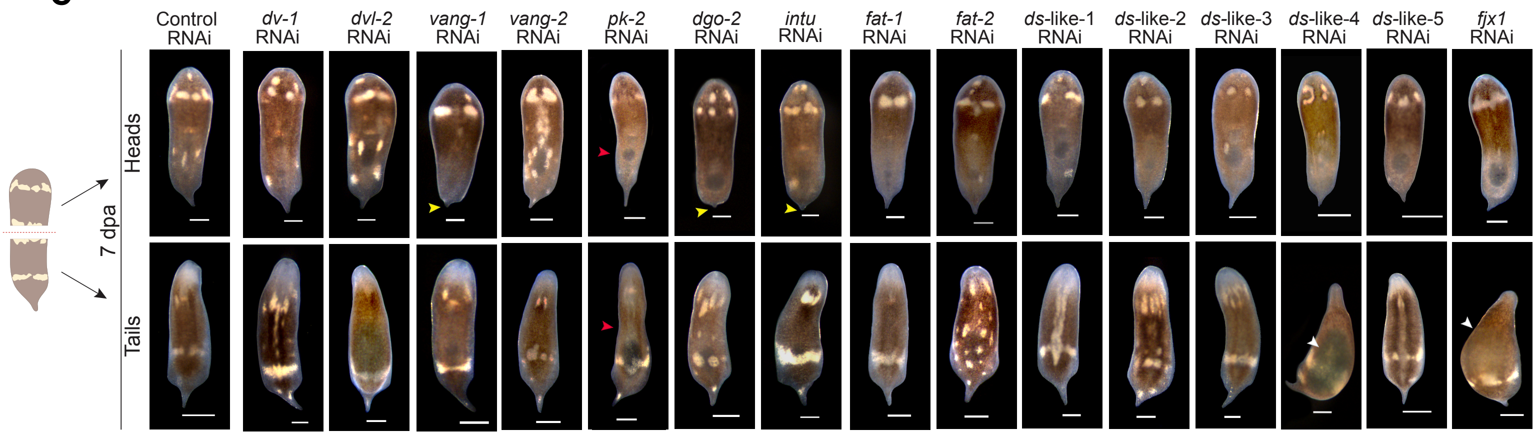


D

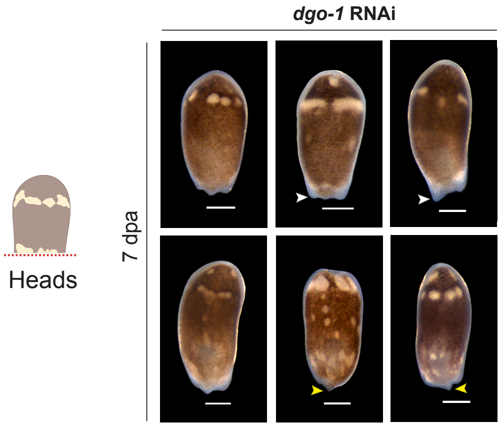

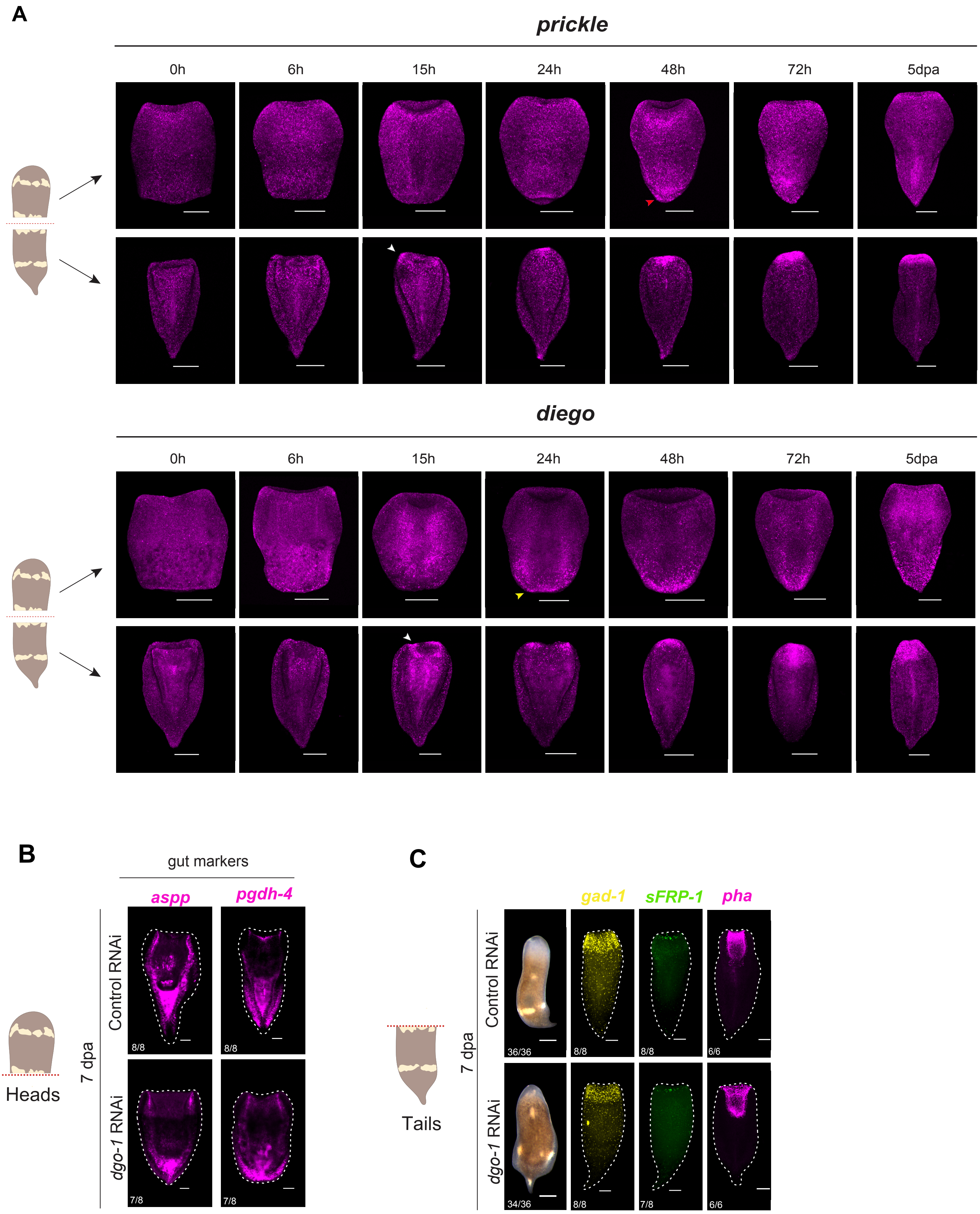

Supp. Figure 3

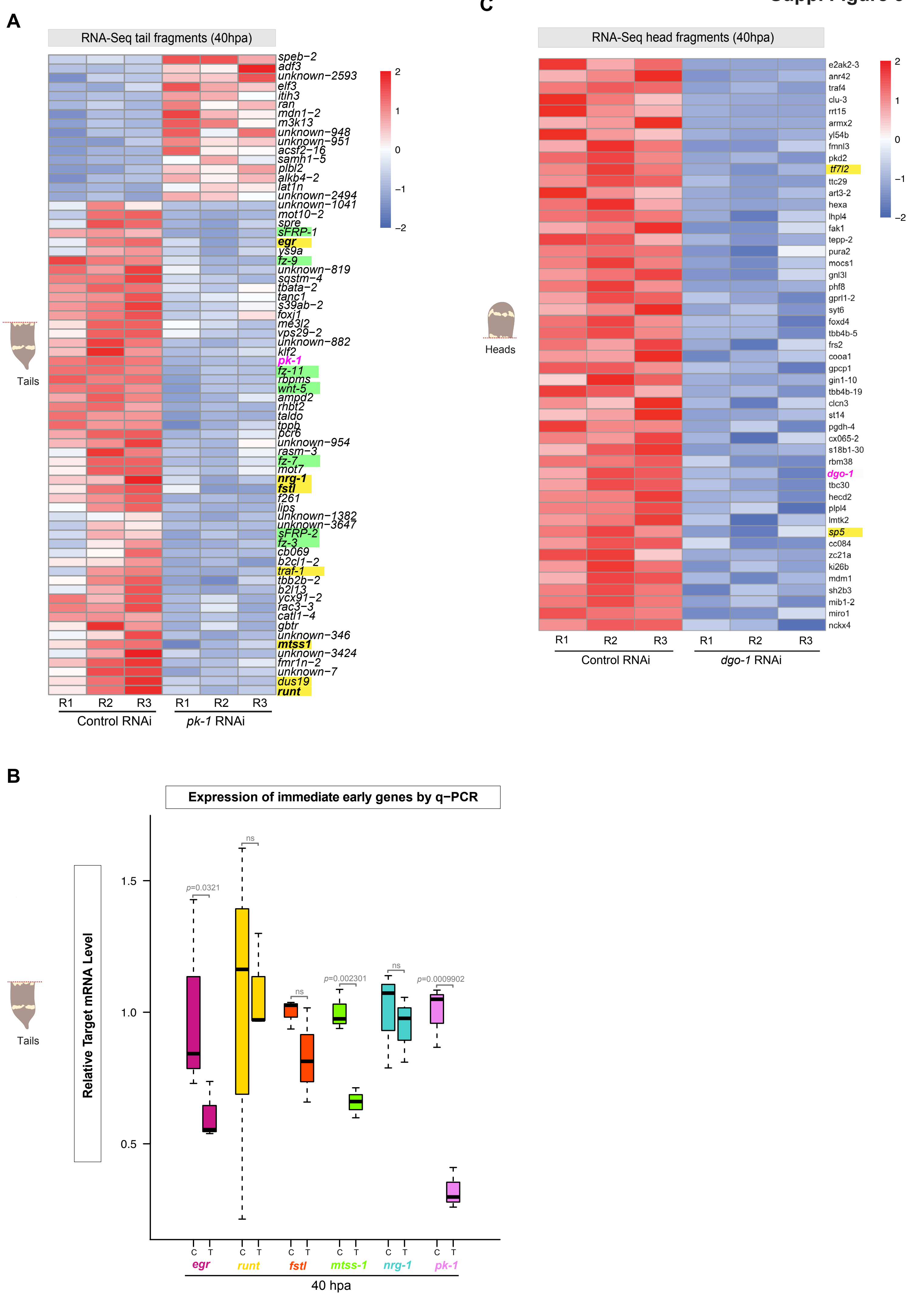

Supp. Figure 3

D

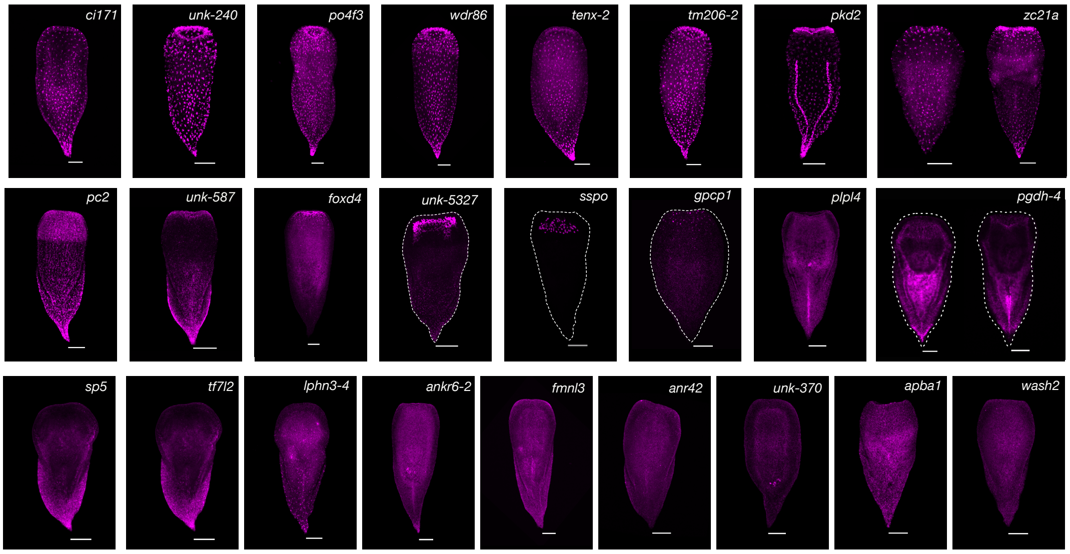


E

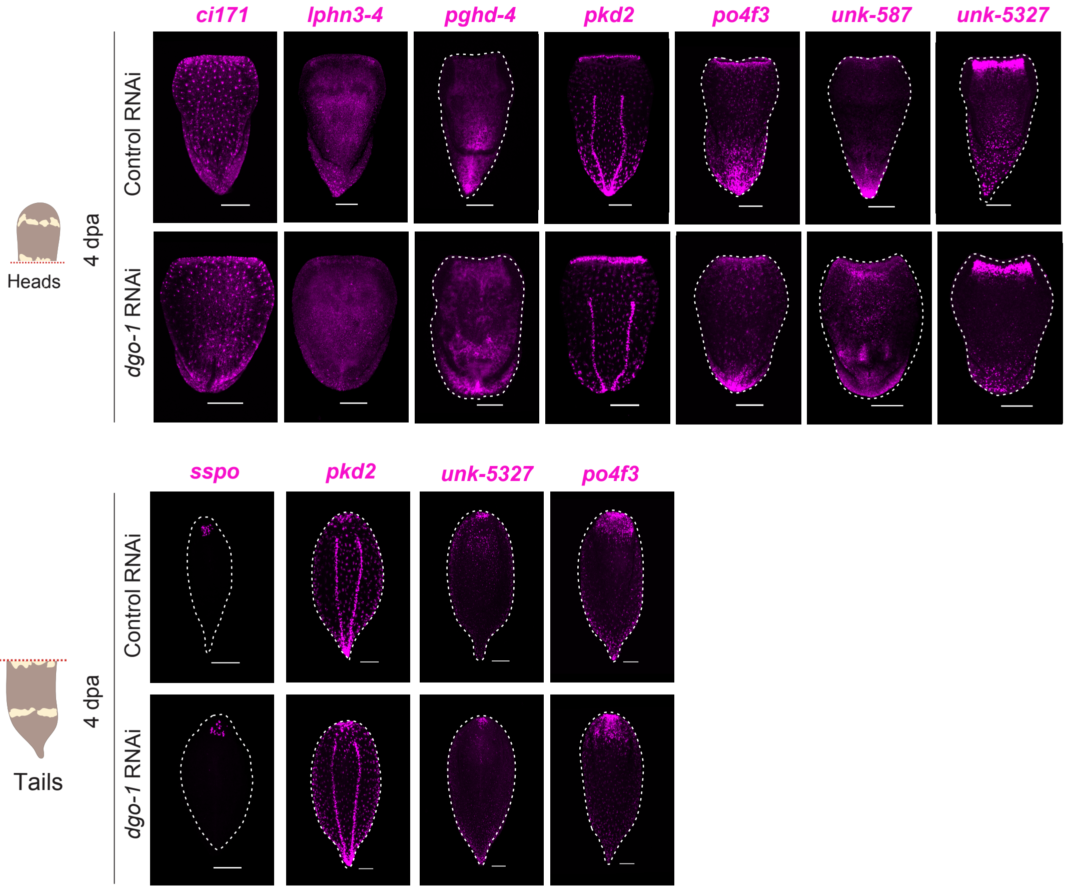

Supp. Figure 4

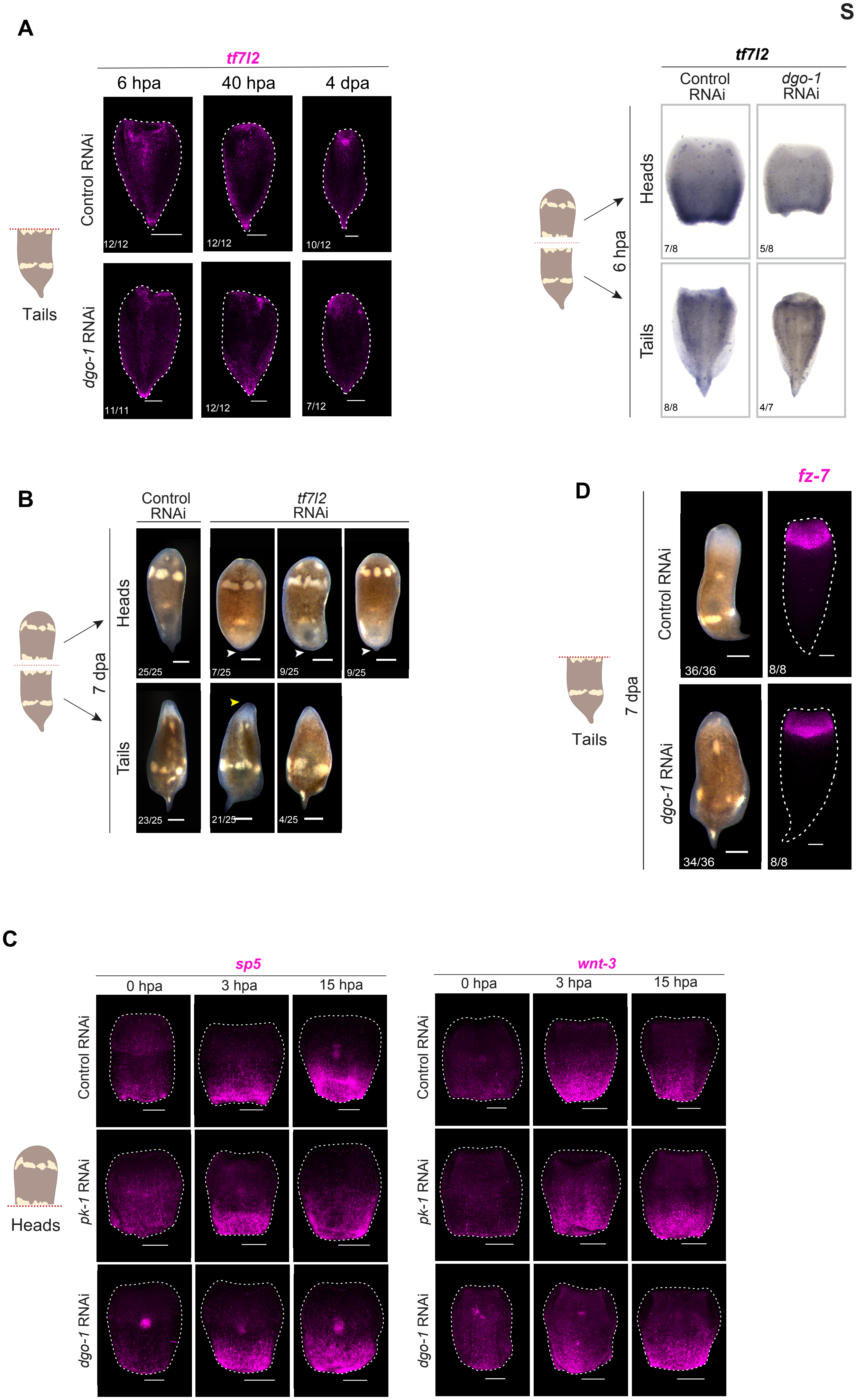

##### Suppl. Figure 5

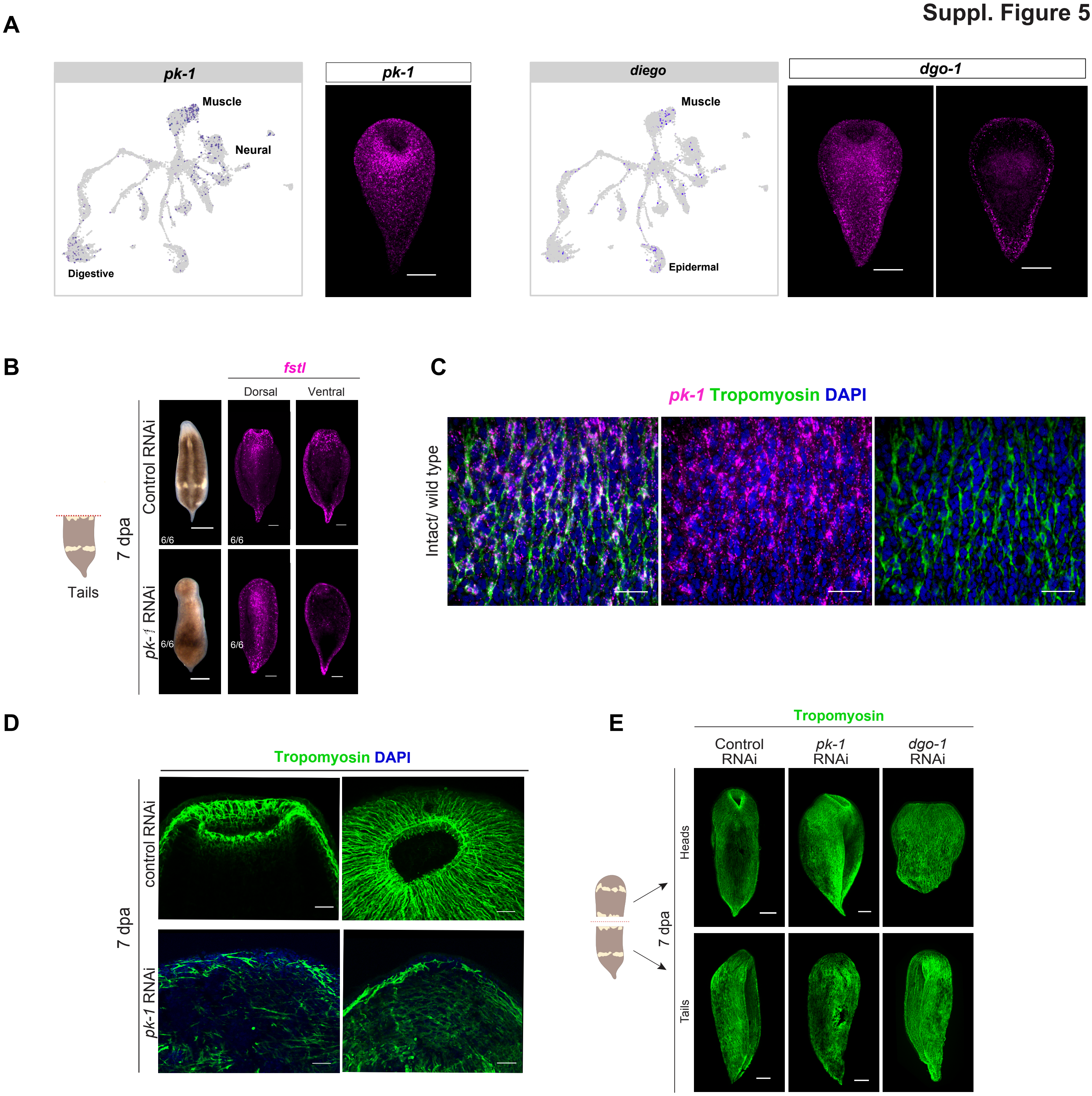
